# Protective effect of estrogen against intervertebral disc degeneration is attenuated by miR-221 through targeting estrogen receptor

**DOI:** 10.1101/184523

**Authors:** Sheng Bin, Yuan Youchao, Liu Xiangyang, Zhang Yi, Liu Hongzhe, Shen Xiongjie, Liu Bin, Chang Lei

## Abstract

Intervertebral disc degeneration (IDD) is a multifactorial disease that associates apoptosis, senescence and calcification of cartilage cells, inflammatory response and alterations in the extracellular matrix. Previous documents imply that estrogen and miR-221 may be involved in IDD. This study further investigated their regulatory mechanisms underlying IDD. Normal and degenerated cartilaginous endplates (CEP) tissues were isolated surgically from juvenile patients with idiopathic scoliosis and adult patients with IDD, respectively. PCR and western blot assays showed decreased aggrecan, Col2A1, TGF-β and estrogen receptorα (ERα) levels in CEP, but increased MMP-3, adamts-5, IL-1β, TNF-α, IL-6 and miR-221 levels. CEP cells were harvested from degenerated CEP tissues and treated with doses of 17β-E2. 17β-E2 increased expression of aggrecan and Col2A1 levels in endplate chondrocytes and secretion of TGF-β, but decreased IL-6 secretion. Moreover, 17β-E2 inhibited the apoptosis and cell-cycle arrest in G0/G1, improving the cell viability. These data indicated estrogen confers protective effect against IDD. However, we found that ERα was a target of miR-221 via luciferase assay. miR-221 up-regulation via the mimics or ERα knockdown attenuated these protective effects conferred by estrogen, while intervention of miR-221 via the inhibitors promoted the protective effects. This study provided novel evidence that estrogen confers protective effects of CEP against IDD, however, up-regulated miR-221 in degenerated CEP decreased the protective effects via targeting ERα, thus it may be an important cause for IDD.

## Introduction

Intervertebral disc degeneration (IDD) is a main cause for low back pain and disability in most regions of the world including China [1]. Intervertebral disc (IVD) consists of the nucleus pulposus (NP), fibrocartilaginous annulus fibrosus (AF) and cartilaginous endplates (CEP). The NP is located in the center of the IVD, which is surrounded and contained by the AF. The CEP is a thin layer of hyaline cartilage that lies above and below the AF serving as main route for the nutrition supply of IVD. The specific structure of IVD helps to transfer and distribute spinal loads between vertebral bodies while allowing mobility. IDD usually begins in the third decade of life and the incidence increases with the age. Some degree of IDD appears to be present in all members over age 50 [2]. IDD is characterized by substantially biomechanical and microstructural changes which reduce the ability of IVD to bear weight and cause a number of pathophysiological features, including lumbago, sciatica, cauda equina symptoms [3]. IVD is rich in proteoglycans held together by a network of type I and/or II collagen fibers. However, IDD exhibits the loss of proteoglycan (PG) and water content in the extracellular matrix, but the increase in matrix-degrading enzymes with the changes of collagen types [4, 5]. These changes contribute to the transition from a gelatinous to a fibrotic NP, which impairs the function of the NP to convert compressive forces into evenly distributed tensile stresses in the surrounding AF [6]. The pathogenesis of IDD is not fully clarified, but it is believed that IDD a multifactorial disease that associates with increased senescence, apoptosis and calcification of nucleus and endplate cartilage cells, inflammatory response, and oxidative stress.

Estrogen probably exerts profound effects on physiological functions of articular cartilage, based on previous documents. Previous study confirmed that menopause is associated with IDD, through investigating 4230 cases of IVD specimens [7]. Thus this study suggests that estrogen decrease may be a risk factor for IDD. Estrogen has been reported to be involved in differentiation of cartilage, although the regulatory effect may be controversial. Study by Maneix et al., [8] shows that 17beta-estradiol (E2) up-regulates cartilage marks including type II collagen gene (COL2A1) and Sox-9 in dedifferentiated articular chondrocytes in osteoarthritis, suggesting the promoting effect of 17β-E2 on chondrocyte redifferentiation. But, a tissue engineering study found that 17β-E2 inhibits chondrogenic differentiation of mesenchymal stem cells via activating G protein-coupled receptor 30 [9]. Estrogen presumably confers different effect on cartilage differentiation, depending on cell types. Estrogen is also involved in the regulation of chondrocyte aging that is commonly observed in the IDD. Estrogen inhibits telomere shortening, thereby retarding senescence of chondrocyte [10].

microRNAs (miRNAs) is an abundant class of small non-coding regulatory RNAs. It commonly functions as an important endogenous silencers of genes through inducing the degradation of target mRNAs. Recently, miRNAs gains substantial attention due to the involvements in various physiological and pathological processes. Report shows that miR-221 is associated with osteogenic differentiation of human annulus fibrosus cells [11]. Endochondral ossifications are common phenomenon in end stages of IVD. AF cells from degenerated discs have a greater tendency for osteogenic differentiation than normal AF cells. According to previous data, overexpression of miR-221 diminishes osteogenic potential of degenerated annulus fibrosus cells in osteogenic differentiation medium, which involves the BMP-Smad pathways [11]. However, miR-221 also blocked the chondrogenic differentiation of human mesenchymal stem cells that are mainly present in bone marrow [12, 13], suggesting that miR-221 may hinder cartilage repair by resident progenitor cells. therefore, the role of miR-221 in IDD remain controversial.

Although estrogen and miR-221 were reported to be involved in IDD, the underlying mechanisms were not elusive. Moreover, previous studies only investigated the functions of estrogen and miR-221 separately, but did not examine the possible relationship between them in IDD. Using the starBase v2.0 prediction software (http://starbase.sysu.edu.cn/index.php), we found that the estrogen receptorα (ERα) was a potential target of miR-221. These study was aimed to find the correlation between estrogen and miR-221 and further detect their regulatory mechanisms in IDD.

## Material & methods

### Isolation of human CEP tissues

Normal and degenerated CEP tissues were isolated surgically from juvenile patients with idiopathic scoliosis and adult patients with IDD, respectively. All patients were informed of the investigational nature of the study. Written informed consent was obtained from them before the experiment. This study was reviewed and approved by the Ethics Committee of Xiangya School of Medicine (Changsha, People’s Republic of China). All tissue samples were subjected to histopathological identification before used for further study.

### Isolation and culture of human endplate chondrocytes (ECs)

Human ECs were obtained from CEP tissues according to previously described method with minor revision [14]. Briefly, the CEP tissues were minced into small pieces (1 mm^3^) and digested with 0.25% trypsin for 30 min at 37°C. After being washed with PBS for three times, the samples were further subject to 0.2% collagenase type II for 4 hours at 37°C. The isolated cells were cultured in complete culture medium (DMEM/F12, Life Technologies, Carlsbad, CA, USA) supplemented with 10% fetal bovine serum (FBS, Invitrogen, Carlsbad, CA, USA) and 1% penicillin/streptomycin (Life Technologies). The second passage of primary EC was used in the whole study.

### Quantitative real-time PCR (qPCR)

Total RNA, including small RNA, was extracted from tissue specimens and ECs using a mirVana miRNA isolation kit (Thermo Fisher Scientific, Waltham, MA, USA). The High Capacity RNA-to-cDNA Master Mix (Life Technologies) was used to synthesize cDNA. Primer sequences were as follow: miR-221 (F) 5′-GGACCTGGCATACAATGT-3′ and (R) 5′-TTTGGCACTAGCACATT-3′; U6 (F) 5′-CTCGCTTCGGCAGCACA-3′ and (R) 5′-AACGCTTCACGAATTTGCGT-3′. TGF-β (F) 5′-AACTATTGCTTCAGCTCCACAGAG-3′ and (R) 5′-AGTTGGATGGTAGCCCTTG-3′; IL-1β (F) 5′-AAACAGATGAAGTGCTCCTTCCAGG-3′ and (R) 5′-TGGAGAACACCACTTGTTGCTCCA-3′; TNF-α (F) 5′-GATGGACTCACCAGGTGAG-3′ and (R) 5′-CTCATGGTGTCCTTTCCAGG-3′; IL-6 (F) 5′-TGCATGACTTCAGCTTTACTCTTTG-3′ and (R) 5′-GGGGAGATAGAGCTTCTCTTTCGTT-3′; β-actin (F) 5′-GTGGGGCGCCCCAGGCACCA-3′ and (R) 5′-CTCCTTAATGTCCGGACGATTC-3′. qPCR was performed using the ABI PRISM 7500 DNA Sequence Detection System (Life Technologies). The relative expression levels were determined using the comparative threshold cycle (2^-ΔΔCT^). The expression level of miR-221 was normalized to that of U6 RNA. The expression levels of other genes were normalized to that of β-actin.

### Western blot assay

Western blot assay was performed to examine expression of indicated protein levels in CEP tissue specimens and ECs. Protein extracts were separated using SDS/polyacrylamide gel electrophoresis and transferred to nitrocellulose membranes. The membranes were hybridized with anti-bodies against aggrecan (ab36861, Abcam, Shanghai, China), Col1A1 (ab34710, Abcam), Col2A1 (ab34712, Abcam), ERα (ab32063, Abcam), Adamts5 (ab41037, Abcam), MMP-3 (ab53015, Abcam) and β-actin (Dilution 1:800, sc-47778, Santa Cruz Biotechnology, Santa Cruz, CA, USA) at 4°C overnight. The primary antibodies were visualized by adding secondary biotin-conjugated antibodies followed by an avidin/biotin/peroxidase complex (Vectastain ABC Elite kit; Vector Laboratories lnc, Burlingame, CA, USA) and substrate (Vector Nova RED, Vectastain).

### Transfections

miR-221 mimics (Sense strand: 5 -AGCUACAUUGUCUGCUGGGUUUC-3 ; Antisense strand: 5 -AACCCAGCAGACAAUGUAGCUUU-3), miR-221 specific inhibitor (5 -GAAACCCAGCAGACAAUGUAGCU-3) and appropriate negative control molecules were purchased from RiboBio Corporation (Guangzhou, China). Small hairpin RNA (shRNA) oligonucleotides against ER (shRNA-ER: 5 -GCTTCAGGCTACCATTATGttcaagagacataATGGTAGCCTGAAGCttttttacgcgt-3) and the corresponding scrambled control oligonucleotides were synthesized from Genephama Biotech (Shanghai, China). These RNA oligoribonucleotides were transfected into ECs using X-tremeGENE siRNA Transfection Reagent (Roche, Germany) according to the manufacturer’s protocol. The final concentration was 125 nM for miRNA mimics, 250 nM for miRNA inhibitors and 60 nM for shRNA. For ectopic expression of ER in ECs, transfection of pEGFP-C1-ER vector (Genephama Biotech) was performed using the Lipofectamine 2000 of Thermo Fisher Scientific, according to the company’s instructions.

### Luciferase assay

The fragments of the 3′ untranslated region (UTR) of ER were amplified by RT-PCR. There are two binding sites at 3′UTR of ER for miR-221 according to the starBase v2.0 prediction software. These two binding sites were mutated separately using site-directed mutagenesis. The wild-type and mutant of 3′UTR of ER were inserted into the pmirGLO vector (XhoI and NotI restriction enzyme sites; Promega, Madison, WI, USA). Vectors were transfected into ECs with miR-221 mimics or not, using Lipofectamine 2000 (Invitrogen). Luciferase and renilla signals were measured 48 h after transfection using the Dual Luciferase Reporter Assay System (Promega). The activity of firefly luciferase was normalized to that of renilla luciferase.

### HE and toluidine blue staining

CEP tissue specimen slices were fixed in 4% paraformaldehyde at 4°C for 30 min. After being rinsed with PBS, slices were stained with hematoxylin and eosin (H&E). ECs were seeded on sterile coverslips in 6-well plates. ECs were initially fixed in 4% paraformaldehyde at 4°C for 30 min. After being rinsed with PBS twice, ECs were stained with 0.1% toluidine blue. Cells were washed extensively and photographed.

### Immunocytochemistry (ICC)

ECs were cultured on coverslips in 6-well plates. Cells were fixed with 4% paraformaldehyde for 10 min at room temperature and with methanol at −20°C for 20 min. Normal goat serum (10%; Hyclone; GE Healthcare Life Sciences) was added to cells for 30 min to block nonspecific binding sites. The fixed cells were immunostained with primary antibodies targeting Col2A1 and SOX9 (Abcam) overnight at 4°C and the Alexa Fluor 488-conjugated secondary antibody (1: 500 dilution, catalog no. SP-9000; ZSGB-BIO, Beijing, China) for 1 h at 37°C. Images were acquired with a high-resolution CoolSNAP™ CCD camera (Photometrics Inc., Tucson, AZ, USA) under the control of a computer using Leica FW4000 software version 1.2 (Leica Microsystems, Ltd., Milton Keynes, UK).

### ELISA assay

Culture supernatants were collected and stored at −20 °C until analysis. The concentration of Col2a1, Aggrecan, Adamts-5 and Mmp-3 in cell culture supernatants was measured using ELISA kits (R&D Systems, Minneapoils, MN, USA) according to the manufacturer’s instructions.

### Cell viability assay

Cells were seeded in 96-well plates. Viable cell numbers were detected by the use of Cell Counting Kit-8 (CCK-8, Dojindo, Kyushu, Japan) following the kit’s instructions.

### Flow cytometry measurement

Apoptosis rate and cell cycle were detected by flow cytometry. Cells were prepared in 6-well plates. Apoptotic incidence was analyzed by the Annexin V-FITC/PI apoptosis detection kit (Beyotime Institute of Biotechnology, Shanghai, China) following the manufacturer’s instructions. Briefly, cells were stained by Annexin V-FITC and PI (propidium iodide) for 15 min in the dark at room temperature. The cells were subjected to a flow cytometer (Beckman Coulter) within 1 h and apoptotic cells were quantified. For cell cycle detection, ECs were fixed in 70% alcohol for 30 minutes on ice. The cells were treated with RNase A (Sigma-Aldrich Co., St Louis, MO, USA) at 37°C and stained with propidium iodide in the dark for 30 minutes. Cell cycle was analyzed by flow cytometry using a FACS-Calibur instrument (BD, Franklin Lakes, NJ, USA).

### Statistical Analysis

Results were presented as means ± standard deviation. Statistical analysis was performed by SPSS 11.0 (SPSS, Chicago, IL, USA). One-way analysis of variance (ANOVA) was used for data analysis, followed by least significant difference test (Fisher test) and the unpaired Student’s t-test was used for comparisons between two means. p values less than 0.05 were considered significant.

## Results

### Difference between the normal and degenerated CEP tissues in the expression of indicated genes and proteins

CEP consists of chondrocytes and a fibril network made from various types of collagen, hyaluronic acid and proteoglycan that enfold the chondrocytes. Previous documents note that proteoglycan is reduced in IDD, while some matrix-degrading enzymes are increased, which changes the biological and physical properties of IVD. Our data showed that aggrecan (the major constituent of proteoglycan) protein level was lower in degenerated CEP tissues than normal CEP tissues (p < 0.05, Figure 1A). Matrix-degrading enzymes, including MMP-3 and Adamts-5, were increased in degenerated CEP tissues compared to the normal tissues (p < 0.05). Col2A1 was decreased in degenerated CEP tissues (p < 0.05), but Col1A1 showed no significant change between normal and degenerated CEP tissues. In addition, this study for the first time found that ERα protein level was decreased in degenerated CEP tissues, compared to normal CEP tissues (p < 0.05).

**Figure 1.**
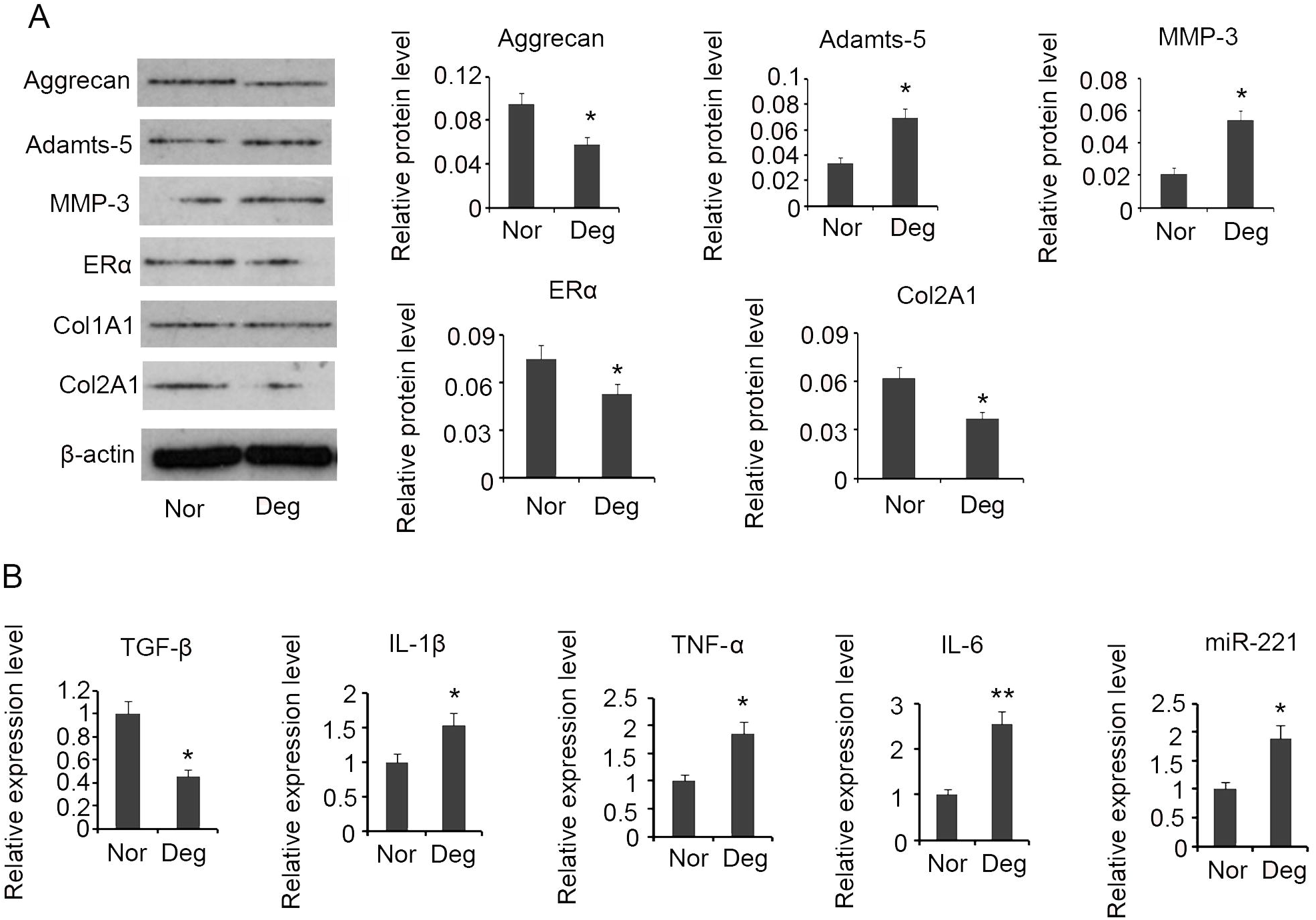
Difference between the normal and degenerated CEP tissues in the expression of indicated genes and proteins. Normal and degenerated cartilaginous endplates tissues were isolated surgically from juvenile patients with idiopathic scoliosis and adult patients with intervertebral disc degeneration, respectively. (A) Western blot assay detected expression of proteins including Aggrecan, Adamts-5, Col2A1, Col1A1, MMP-3 and estrogen receptor α. (B) PCR assay detected TGF-β, IL-1β, TNF-α, IL-6 and miR-221 expression. *P<0.05 and **P<0.01 *vs.* normal tissues. Nor: normal cartilaginous endplates tissues; Deg: degenerated cartilaginous endplates tissues

Previous studies reveals that some cytokines, like inflammatory cytokines, probably involved in pathogenesis of IDD. Using RT-PCR, this study found TGF-β (p < 0.05, Figure 1B) was reduced in degenerated CEP tissues compared to the normal tissues, while IL-1β (p < 0.05), TNF-α (p < 0.05) and IL-6 (p < 0.01) were increased in degenerated CEP tissues. Moreover, we found that miR-221 (p < 0.05) was notably increased in degenerated CEP tissues as well.

### Regulatory effect of estrogen on chondrocytes from degenerated CEP tissues

To understand the regulatory effect of estrogen on degenerated CEP, we performed an in vitro study in endplate chondrocytes isolated from degenerated CEP tissues. Figure 2A shows the HE staining of degenerated CEP tissues. Toluidine blue staining showed that the isolated endplate chondrocytes were polygonal or spindle-shaped (Figure 2B). Col2A1 and SOX9 are important markers for chondrocytes. Immunofluorescence staining for Col2A1 and SOX9 verified their distribution in the cytoplasm (Figure 2C).

**Figure 2.**
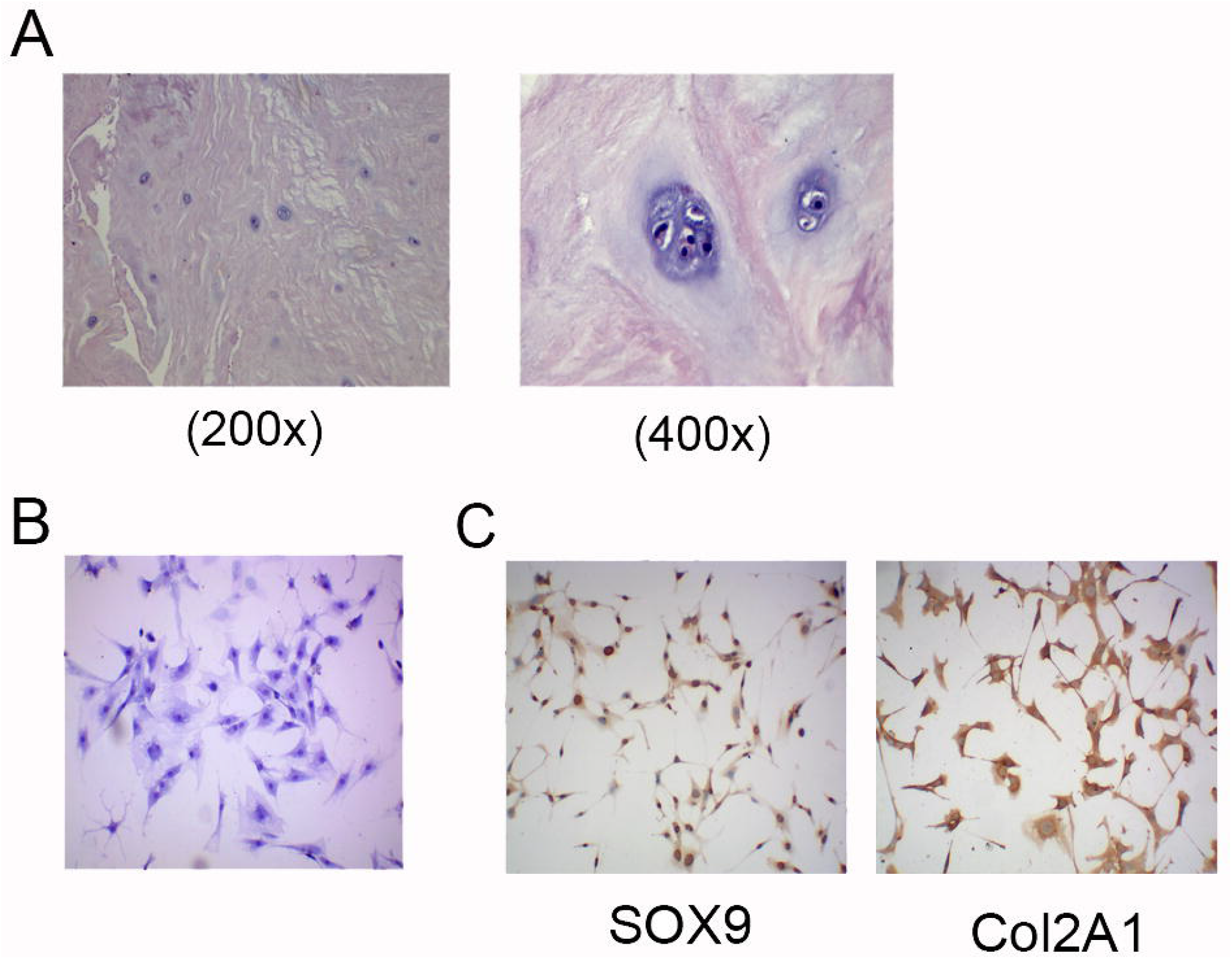
Isolation of chondrocytes from degenerated CEP tissues. (A) HE staining of degenerated CEP tissues. (B) Toluidine blue staining of chondrocytes isolated from degenerated CEP tissues (400×). (C) Identification of the isolated chondrocytes via ICC (400×).

Isolated endplate chondrocytes were exposed to doses of 17β-E2 for 48h. Western blotting revealed that expression of aggrecan level was increased by 17β-E2 dose-dependently. 10^−8^ (p < 0.05), 10^−7^ (p < 0.01) and 10^−6^ (p < 0.01) mol/L 17β-E2 triggered significant increase in aggrecan (Figure 3A). 17β-E2 at concentrations of 10^−8^ (p < 0.05), 10^−7^ (p < 0.05) and 10^−6^ (p < 0.05) mol/L also increased Col2A1 level in endplate chondrocytes. According to data of ELISA assay, IL-6 concentration in cell medium was reduced by 17β-E2 in a dose-dependent manner (Figure 3B). Notable reduction in IL-6 concentration was observed with 10^−7^ and 10^−6^ mol/L 17β-E2. TGF-β concentration was conversely increased by 17β-E2 at dosages of 10^−8^ (p < 0.05), 10^−7^ (p < 0.01) and 10^−6^ (p < 0.01) mol/L (Figure 3C). Treatment with 17β-E2 did not significantly affect IL-1β and TNF-α concentrations in cell medium (data not shown).

**Figure 3.**
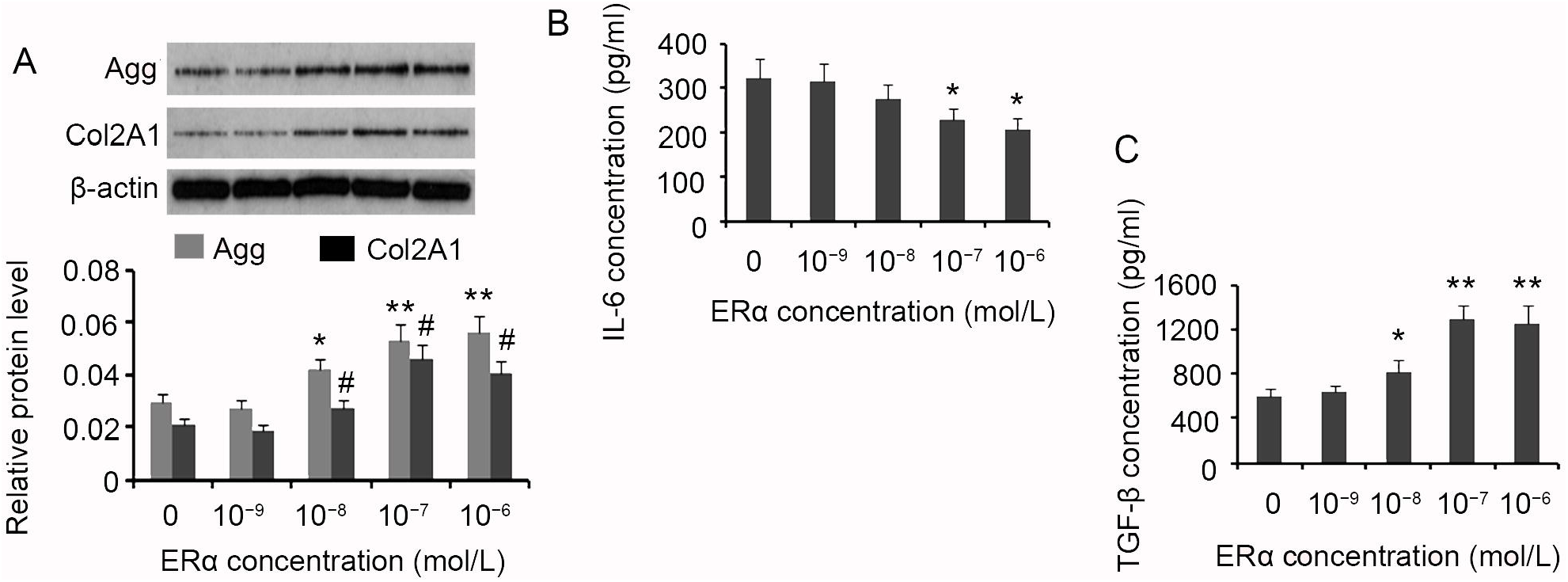
Influence of 17β-E2 on expression of aggrecan and Col2A1 in the ECs and on the secretion of TGF-β and IL-6. Isolated ECs were exposed to doses of 17β-E2 for 48h. (A) Western blotting detected the expression of aggrecan and Col2A1 in the ECs. ELISA assay evaluated (B) TGF-β and (C) IL-6 concentrations in the cell medium. *P<0.05, **P<0.01, ^#^P<0.05 and ^##^P<0.01 *vs.* control. Agg: aggrecan.

10^−7^ mol/L 17β-E2 increased cell viability of endplate chondrocytes, compared to control group (p < 0.05, Figure 4A). The apoptosis rate was decreased by 10^−7^ and 10^−6^ mol/L 17β-E2, as reflected by flow cytometry measurement (Figure 4B). Treatment with 10^−7^ mol/L 17β-E2 decreased cell percentage in G0/G1 cell phase (p < 0.05), but increased cell percentage in S phase (p < 0.05). 10^−6^ mol/L 17β-E2 increased cell percentage in G2/M phase (Figure 4C).

**Figure 4.**
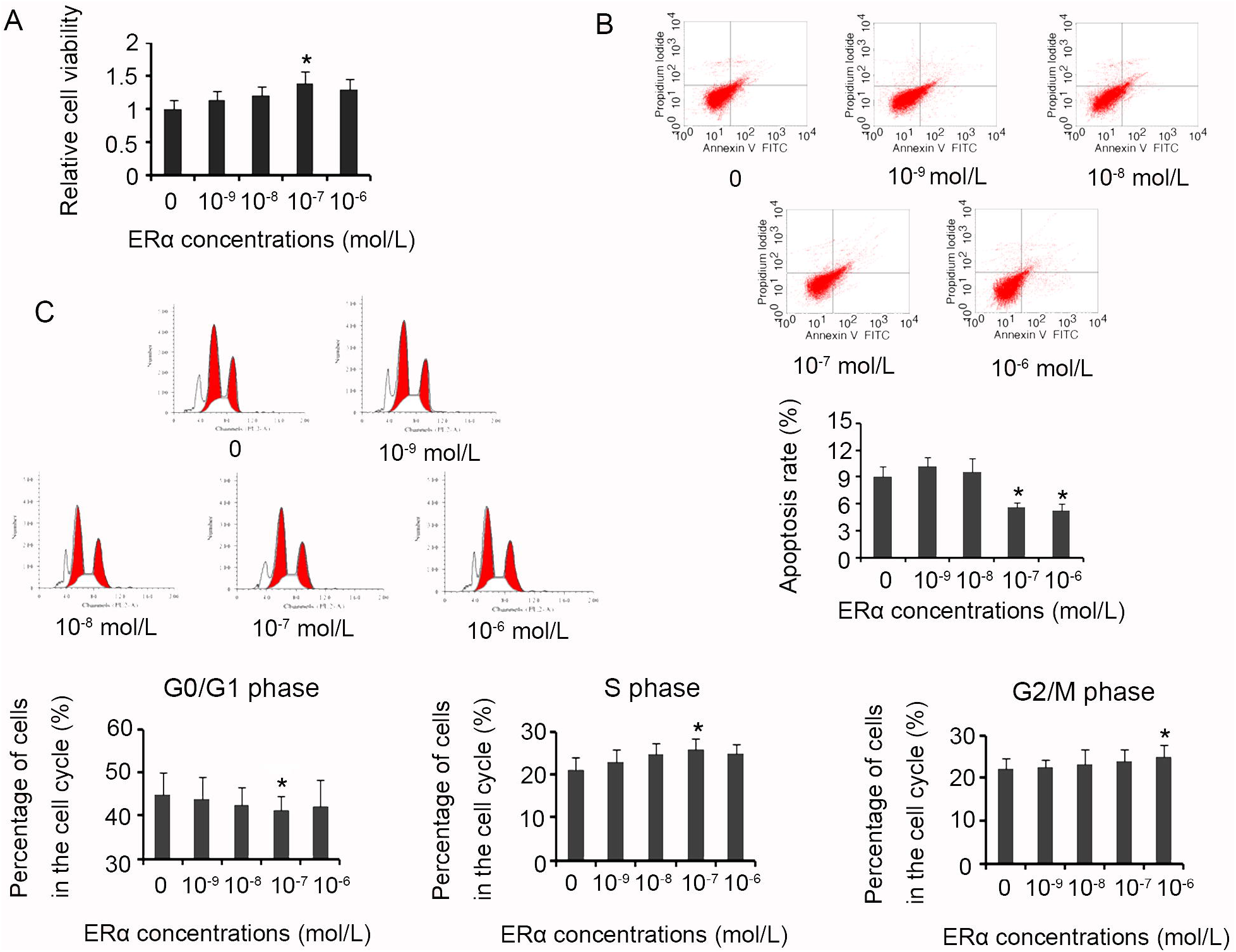
Influence of 17β-E2 on cell proliferative characters of ECs. Isolated ECs were exposed to doses of 17β-E2 for 48h, followed by evaluation of (A) cell viability, (B) apoptosis rate and (C) cell cycle. *P<0.05 and **P<0.01 *vs.* control.

### miR-221 inhibits the effect of estrogen in endplate chondrocytes via targeting ERα

Although miR-221 and estrogen were regarded as candidate modulators implicated in IDD pathogenesis, their correlation has never been reported. We found that ERα might be a target of miR-221 using starBase v2.0 prediction software. To further corroborate this finding, we performed luciferase assay. Reporter vectors containing confragments of the 3′UTR of ERα (RV-ERα) showed decreased luciferase activity in endplate chondrocytes, compared to empty vectors (p < 0.05 Figure 5A). Transfection with miR-221 mimics further decreased luciferase activity of RV-ERα (p < 0.01). There were two potential binding sites in 3′UTR of ERα by miR-221, according to starBase v2.0 prediction software. Mutation at both sites in RV-ERα partly restored luciferase activity (p < 0.05 vs. with-type RV-ERα). These data indicated that ERα is a target of miR-221 in endplate chondrocytes.

**Figure 5.**
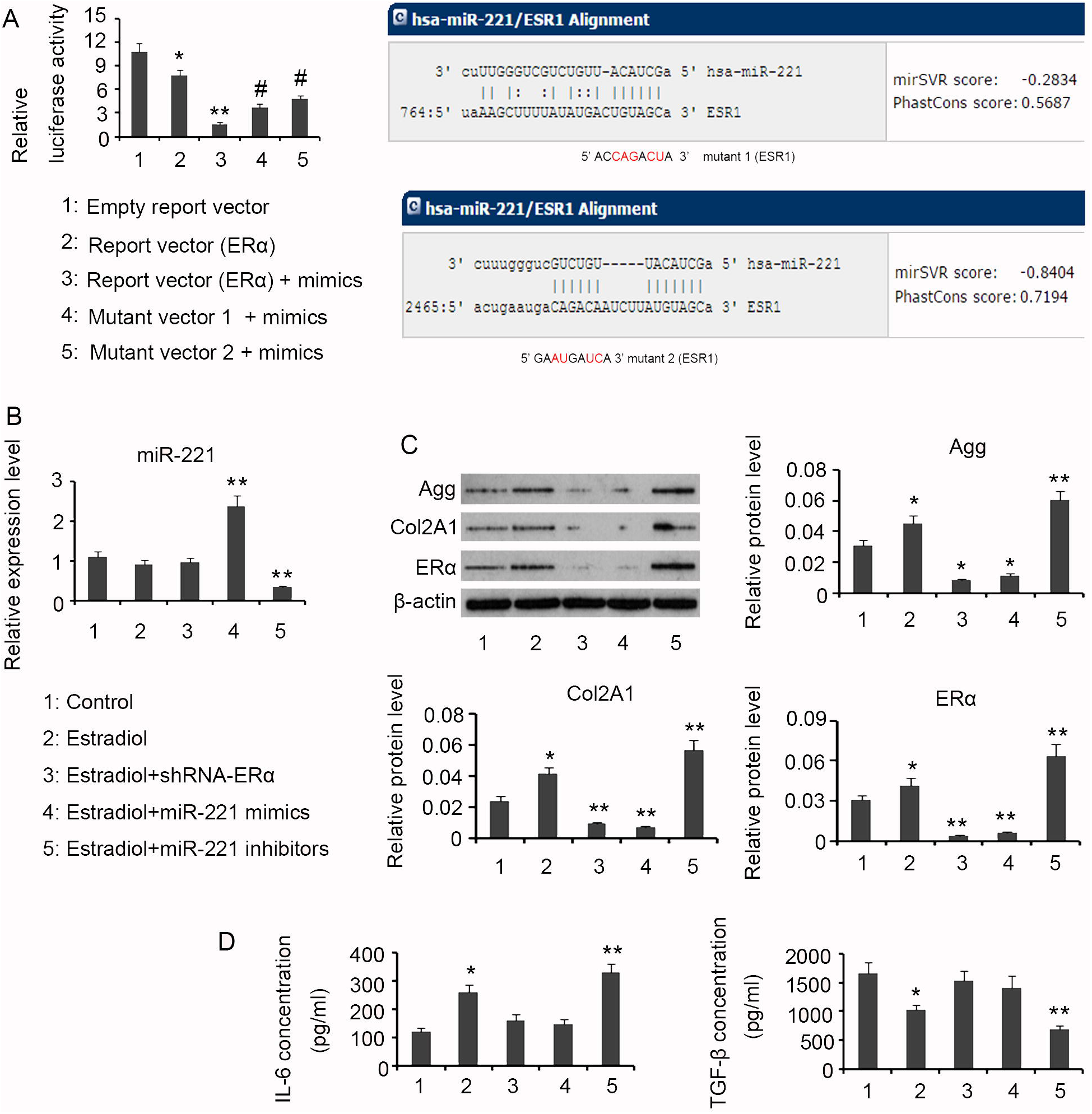
miR-221 targeting ERα influences the downstream effecters. (A) Luciferase assay identified that ERα is a target of miR-221. (B) PCR assay detected miR-221 expression. (C) Western blotting assessed expression of proteins including aggrecan, Col2A1 and estrogen receptor α. (D) ELISA assay evaluated TGF-β and IL-6 concentrations in the cell medium. *P<0.05 and **P<0.01 *vs.* control; ^#^P<0.05 and ^##^P<0.01 *vs.* Reporter vector (ERα) + mimics group. Agg: aggrecan.

To determine the regulatory effects of estrogen in endplate chondrocytes is influenced by miR-221, we undertaken a series of transfection assays. Treatment with 10^−7^ mol/L 17β-E2 alone or in combination with ERα knockdown did not affect miR-221 expression. Transfection with miR-221 mimics and inhibitors respectively increased and decreased miR-221 expression (p < 0.01, Figure 5B). Western blot assay showed that ERα knockdown reversed the increase in aggrecan (p < 0.05), Col2A1 (p < 0.01) and ERα (p < 0.01) by 17β-E2 (Figure 5C). Combination of 17β-E2 treatment and miR-221 overexpression reduced expression of aggrecan (p < 0.05), Col2A1 (p < 0.01) and ERα (p < 0.01) levels. Interruption of miR-221 via inhibitors enhanced the promoting effect of 17β-E2 on expression of aggrecan (p < 0.01), Col2A1 (p < 0.01) and ERα (p < 0.01) levels. Silencing ERα or forcing miR-221 expression abrogated the regulatory effect of 17β-E2 on IL-6 and TGF-β secretion, as indicated by ELISA assay (Figure 5C). Inhibition of miR-221 expression promoted the increase in IL-6 in medium by 17β-E2 (p < 0.01). 17β-E2-triggered reduction in TGF-β in medium was enhanced with miR-221 down-regulation (p < 0.01).

ERα knockdown reversed cell viability that was increased by 17β-E2 (p < 0.05), compared to control (Figure 6A). Down-regulating miR-221 viability further increased cell viability promoted by 17β-E2 (p < 0.05). 17β-E2 inhibited the apoptosis rate of endplate chondrocytes, which was conversely increased by ERα knockdown and miR-221 overexpression (p < 0.05, Figure 6B). miR-221 down-regulation promoted apoptosis rated reduction induced by 17β-E2 (p < 0.05). Analysis of cell cycle showed that cell percentage in G0/G1 phase decreased by treatment with 10^−7^ mol/L 17β-E2 alone or in combination with miR-221 depletion (Figure 6C). Combination of 17β-E2 treatment and miR-221 overexpression increased the cell percentage in G0/G1 phase (p < 0.05). Cell percentage in S phase was increased by 17β-E2 alone or in combination with miR-221 deletion (p < 0.05). However, silencing ERα or forcing miR-221 expression decreased cell percentage in S phase in the presence of 17β-E2 (p < 0.05). miR-221 overexpression decreased cell percentage in G2/M phase in despite of 17β-E2 treatment (p < 0.05). Lost of miR-221 increased cell percentage in G2/M phase in the presence of 17β-E2 (p < 0.05).

**Figure 6.**
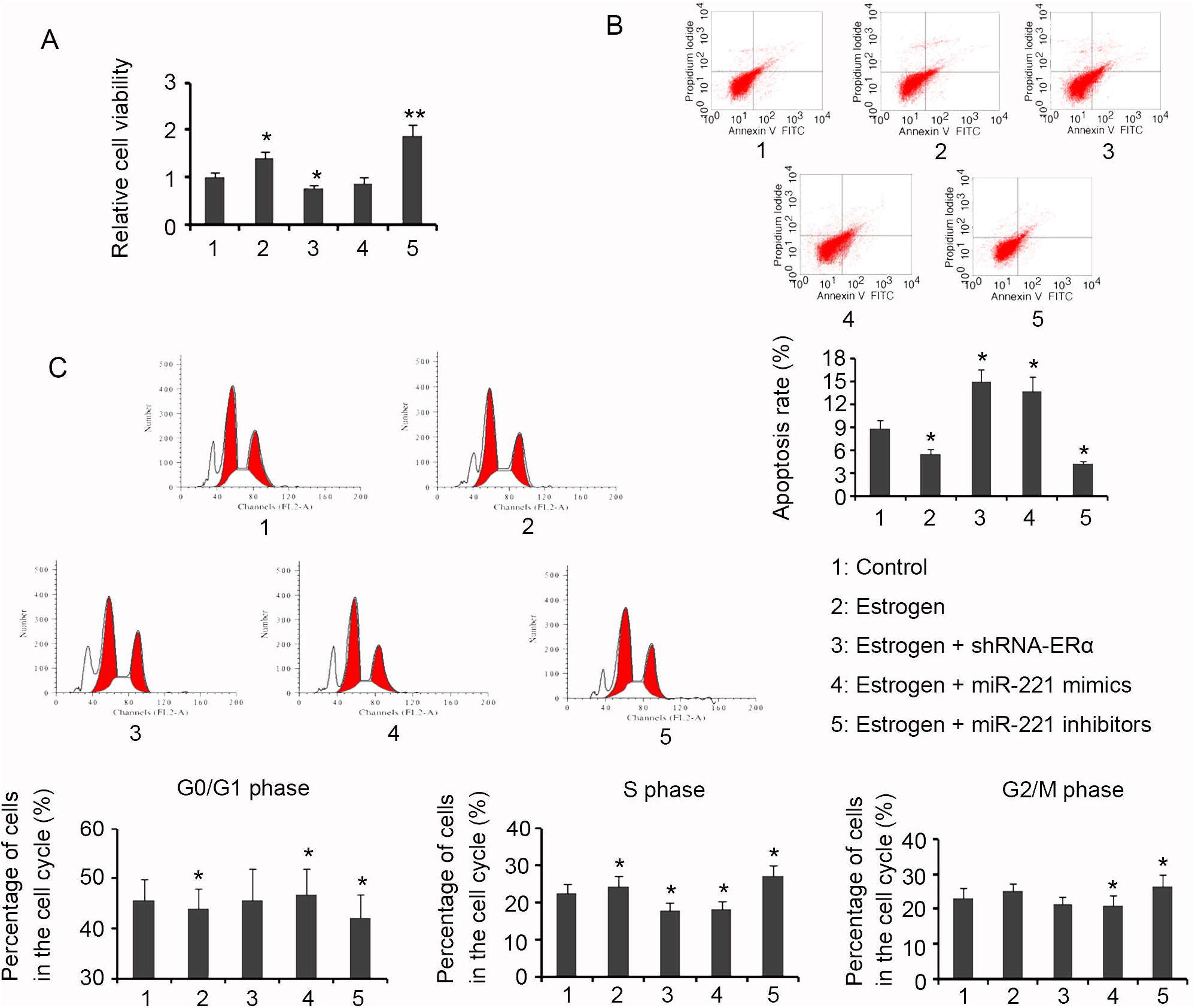
Influence of miR-221 on cell proliferative characters of ECs. ECs were treated with 17β-E2 and/or subjected to transfection assays, followed by evaluation of (A) cell viability, (B) apoptosis rate and (C) cell cycle. *P<0.05 and **P<0.01 *vs.* control.

## Discussion

IDD is a multifactorial disease which associates apoptosis, senescence and calcification of nucleus and endplate cartilage cells, inflammatory response, and oxidative stress [1–3]. Degenerated intervertebral disc shows substantially biomechanical and microstructural changes compared with normal intervertebral disc, thus loses the ability to distribute spinal loads and causes painful symptoms and diseases. IDD is characterized by reduction in proteoglycan in extracellular matrix probably due to increased abundance and activity of matrix-degrading enzymes [1–3]. This study compared degenerated CEP tissues with normal CEP tissues, exhibiting decreased aggrecan and Col2A1 with increased expression of MMP-3 and adamts-5 levels in the degenerated CEP tissues. Previous study revealed that menopause is an import risk factor for IDD, which implies that estrogen probably exerts protective effect against degeneration of cartilage. But the mechanisms by which estrogen antagonizes cartilage degeneration are not fully understood. This study isolated chondrocytes from degenerated CEP tissues and cultured with doses of 17β-E2. Results showed that 17β-E2 promoted expression of aggrecan and Col2A1 of endplate chondrocytes, indicating that 17β-E2 helps to maintain cartilaginous phenotype.

Our data further showed that 17β-E2 stimulated TGF-β secretion from endplate chondrocytes, whereas suppressed their production of IL-6. TGF-β is a multifunctional factor that plays critical role in maintaining intervertebral disc homeostasis [4]. It acts upstream of aggrecan and connective tissue growth factor, regulating their synthesis and the formation of extracellular matrix of cartilage. Extensive evidence shows that TGF-β drives the differentiation of human mesenchymal stem cells toward NF-like cells under simulated microgravity conditions, supports extracellular matrix formation by AF cells, and prevents abnormal calcification of CEP through increasing Ank expression [15, 16]. TGF-β inhibits NF-κB and MAPK pathways, thereby serving as a strong immune suppressor in IVD [17]. Inflammatory immune responses are critically involved in the pathogenesis of IDD. Inflammatory cytokines, such as TNF-α, IL-1β and IL-6, were up-regulated in IDD, as observed in previous studies and the present study [5]. Inflammatory cytokines lead to up-regulation of matrix-degrading enzymes, like MMP-3, and facilitate apoptosis and senescence of chondrocytes. In this study, 17β-E2 suppressed production of IL-6 by endplate chondrocytes, but did not affect the secretion of TNF-α and IL-1β.

Notable reduction in the number of viable and functional cells in the microenvironment of IVD is also an important cause for IDD [18]. Massive cell loss ultimately leads to deterioration in quality or function of tissues or organs and possible failure of vital organs, which is a common core feature of most degenerative diseases. Increased cell death in IDD associates excessive apoptosis and senescence [19]. Recent researches indicate that signaling pathways that mediate apoptosis and senescence was abnormally activated in response to numerous factors, such as nutrient depletion, environment influences, physical impairment, inflammatory stimulation and oxidative stress [19]. However, protection against apoptosis and senescence of chondrocytes ameliorates IDD in vivo [19]. This study showed that 17β-E2 increased viability of chondrocytes, reducing the apoptosis rate. Cell-cycle analysis indicated that 17β-E2 inhibits cell cycle arrest in G0/G1 phase, which can accelerate cell proliferation, thus maintaining functional and morphological integrity of IVD.

Although the estrogen conferred many positive effects against IDD, these effects were attenuated by the up-regulated miR-221. This study identified that miR-221 binds to 3’UTR of ERα at two binding sites, resulting in the interruption of ERα expression. miR-221 was up-regulated in degenerated CEP tissues compared to the normal tissues. It is probably an important cause for ERα reduction in degenerated CEP tissues. In the chondrocytes isolated from degenerated CEP tissues, we found that loss of ERα impaired protective effects of estrogen against IDD, with the consequence that TGF-β expression was decreased, while IL-6 was increased. The cell viability was decreased in accompany with elevated apoptosis rate. CEP is pivotal for IVD, as it is the only source for NP and AF cells acquiring nutrients and oxygen. Study discovers that nutrients and oxygen tension within the disc are usually reduced as a result of the dysfunction of CEP [20]. Therefore, impaired function of CEP exacerbate metabolic imbalance of the center of the IVD, ultimately triggering IDD.

This study provided novel evidence that estrogen confers protective effects against IDD. Estrogen increased expression of aggrecan and Col2A1 levels in endplate chondrocytes and secretion of TGF-β; It decreased IL-6 secretion from endplate chondrocytes and the apoptosis. However, due to the up-regulation of miR-221, ERα was down-regulated in degenerated CEP tissues, and thus the protective effects of estrogen were profoundly impaired. These data suggest that miR-221 may be a promising target to enhance protective effect of estrogen against IDD.

## Acknowledgement

This study is funded by the General Project of Hunan Health and Family Planning Commission (B2016008).

## References

1. Wu X, Song Y, Liu W, Wang K, Gao Y, Li S, Duan Z, Shao Z, Yang S, Yang C. IAPP modulates cellular autophagy, apoptosis, and extracellular matrix metabolism in human intervertebral disc cells. Cell Death Discov. 2017 Jan 30;3:16107.

2. Jin H, Shen J, Wang B, Wang M, Shu B, Chen D. TGF-β signaling plays an essential role in the growth and maintenance of intervertebral disc tissue. FEBS Lett. 2011 Apr 20;585(8):1209–15.

3. Li YF, Tang XZ, Liang CG, Hui YM, Ji YH, Xu W, Qiu W, Cheng LM. Role of growth differentiation factor-5 and bone morphogenetic protein type II receptor in the development of lumbar intervertebral disc degeneration. Int J Clin Exp Pathol. 2015 Jan 1;8(1):719–26.

4. Bian Q, Ma L, Jain A, Crane JL, Kebaish K, Wan M, Zhang Z, Edward Guo X, Sponseller PD, Séguin CA, Riley LH, Wang Y, Cao X. Mechanosignaling activation of TGFβ maintains intervertebral disc homeostasis. Bone Res. 2017 Mar 21;5:17008.

5. Qin C, Zhang B, Zhang L, Zhang Z, Wang L, Tang L, Li S, Yang Y, Yang F, Zhang P, Yang B. MyD88-dependent Toll-like receptor 4 signal pathway in intervertebral disc degeneration. Exp Ther Med. 2016 Aug;12(2):611–618.

6. Arkesteijn IT, Smolders LA, Spillekom S, Riemers FM, Potier E, Meij BP, Ito K, Tryfonidou MA. Effect of coculturing canine notochordal, nucleus pulposus and mesenchymal stromal cells for intervertebral disc regeneration. Arthritis Res Ther. 2015 Mar 14;17:60.

7. Lou C, Chen HL, Feng XZ, Xiang GH, Zhu SP, Tian NF, Jin YL, Fang MQ, Wang C, Xu HZ. Menopause is associated with lumbar disc degeneration: a review of 4230 intervertebral discs. Climacteric. 2014 Dec;17(6):700–4.

8. Maneix L, Servent A, Porée B, Ollitrault D, Branly T, Bigot N, Boujrad N, Flouriot G, Demoor M, Boumediene K, Moslemi S, Galéra P. Up-regulation of type II collagen gene by 17β-estradiol in articular chondrocytes involves Sp1/3, Sox-9, and estrogen receptor α. J Mol Med (Berl). 2014 Nov;92(11): 1179–200.

9. Jenei-Lanzl Z, Straub RH, Dienstknecht T, Huber M, Hager M, Grässel S, Kujat R, Angele MK, Nerlich M, Angele P. Estradiol inhibits chondrogenic differentiation of mesenchymal stem cells via nonclassic signaling. Arthritis Rheum. 2010 Apr;62(4):1088–96.

10. Breu A, Sprinzing B, Merkl K, Bechmann V, Kujat R, Jenei-Lanzl Z, Prantl L, Angele P. Estrogen reduces cellular aging in human mesenchymal stem cells and chondrocytes. J Orthop Res. 2011 Oct;29(10):1563–71.

11. Yeh CH, Jin L, Shen F, Balian G, Li XJ. miR-221 attenuates the osteogenic differentiation of human annulus fibrosus cells. Spine J. 2016 Jul;16(7):896–904.

12. Lolli A, Narcisi R, Lambertini E, Penolazzi L, Angelozzi M, Kops N, Gasparini S, van Osch GJ. Silencing of antichondrogenic MicroRNA-221 in human mesenchymal stem cells promotes cartilage repair in vivo. Stem Cells. 2016;34(7):1801–11.

13. Lolli A, Lambertini E, Penolazzi L, Angelozzi M, Morganti C, Franceschetti T, Pelucchi S, Gambari R, Piva R. Pro-chondrogenic effect of miR-221 and slug depletion in human MSCs. Stem Cell Rev. 2014 Dec;10(6):841–55.

14. Mern DS, Tschugg A, Hartmann S, Thomé C. Self-complementary adeno-associated virus serotype 6 mediated knockdown of ADAMTS4 induces longterm and effective enhancement of aggrecan in degenerative human nucleus pulposus cells: A new therapeutic approach for intervertebral disc disorders. PLoS One. 2017 Feb 16;12(2):e0172181.

15. Hegewald AA, Zouhair S, Endres M, Cabraja M, Woiciechowsky C, Thomé C, Kaps C. Towards biological anulus repair: TGF-β3, FGF-2 and human serum support matrix formation by human anulus fibrosus cells. Tissue Cell. 2013 Feb;45(1):68–76.

16. Colombier P, Clouet J, Boyer C, Ruel M, Bonin G, Lesoeur J, Moreau A, Fellah BH, Weiss P, Lescaudron L, Camus A, Guicheux J. TGF-β1 and GDF5 act synergistically to drive the differentiation of human adipose stromal cells toward nucleus pulposus-like cells. Stem Cells. 2016 Mar;34(3):653–67.

17. Cai F, Zhu L, Wang F, Shi R, Xie XH, Hong X, Wang XH, Wu XT. The paracrine effect of degenerated disc cells on healthy human nucleus pulposus cells is mediated by MAPK and NF-κB pathways and can be reduced by TGF-β 1. DNA Cell Biol. 2017 Feb;36(2):143–158.

18. Gruber HE, Norton HJ, Hanley EN Jr. Anti-apoptotic effects of IGF-1 and PDGF on human intervertebral disc cells in vitro. Spine. 2000 Sep 1;25(17):2153–7.

19. Chen D, Xia D, Pan Z, et al. Metformin protects against apoptosis and senescence in nucleus pulposus cells and ameliorates disc degeneration in vivo. Cell Death Dis. 2016 Oct 27;7(10):e2441.

20. Shirazi-Adl A, Taheri M, Urban JP. Analysis of cell viability in intervertebral disc: Effect of endplate permeability on cell population. J Biomech. 2010 May 7;43(7):1330–6.

